# Parental immune priming reshapes offspring growth, metabolism, and thermal tolerance in the Pacific Oyster

**DOI:** 10.64898/2025.12.10.693539

**Authors:** Madeline Baird, Ariana S. Huffmyer, Noah Ozguner, Samuel J. White, Valentina Valenzuela, Steven B. Roberts

## Abstract

Pacific Oysters (*Magallana*/*Crassostrea gig*as) are marine bivalves that are widely cultivated but increasingly experience summer mortality due to interacting stressors. Two major concerns are (1) the rising severity and frequency of marine heat waves and (2) disease outbreaks (e.g., OsHV-1). It is critical to understand how these interacting stressors influence oyster resilience and how stress memory is passed down from parent to offspring. Specifically, it is not known how parental immune stress impacts offspring tolerance to other stressors, including heat stress. Therefore, we tested the effects of parental immune challenge on offspring performance under elevated temperature. Broodstock were exposed to a Poly(I:C) immune challenge, then their offspring were reared to the seed stage and assessed for survival, growth, and metabolic responses under thermal stress in the lab. We found that parental immune priming elicited metabolic flexibility in offspring, which may underlie altered stress tolerance. Offspring of immune-challenged parents were 6.5% larger by 236 days post-fertilization at the end of the study. Under elevated temperatures, offspring from treated parents had 35% lower mortality than controls at 40°C, but this was reversed at 42°C with 17% higher mortality compared to controls, suggesting thermal limits to priming benefits. Metabolic assays further revealed that at moderately elevated temperature (36°C), primed offspring had 43% higher metabolic activity, whereas at higher temperature (40°C), they exhibited 30% lower metabolic activity than controls. Offspring of immune primed parents differentially expressed heat shock (HSP70, HSP90) and immune genes (viperin) under acute stress, suggesting molecular mechanisms underlying immune priming effects. This pattern indicates that parental immune challenge may influence offspring metabolic flexibility, potentially enhancing thermal tolerance through an increased capacity for metabolic depression at extreme temperatures. Together, the results of this study highlight cross-generational links between immune priming and thermal tolerance.

## Introduction

Pacific oysters (*Magallana gigas*, formerly *Crassostrea gigas* (Thunberg, 1793); (Salvi and Mariottini, 2016)) are bivalve mollusks that play a vital role in coastal ecosystems and global aquaculture. They provide key ecosystem services, including water filtration and habitat structure, while also supporting local economies through their high commercial value (Zhang et al., 2012). Due to their rapid growth and adaptability, *M. gigas* is widely farmed across the United States, particularly in the state of Washington, contributing almost $1 million in production in 2013 (WSG, 2015).

Under climate change and local anthropogenic stressors, Pacific oysters are highly vulnerable to mortality events (Heo et al., 2023; Malham et al., 2009). In both farmed and natural settings, oysters experience “sudden unusual mortality syndrome” (SUMS) events, typically in late spring and summer, driven by combinations of stressors including elevated temperatures, hypoxia, and pathogen outbreaks. SUMS events can lead to 30-90% mortality occurring in short periods with no clear known cause (Ben-Horin et al., 2025; Bodenstein et al., 2023; Guévélou et al., 2019; Matt et al., 2020). Increasing environmental stress resulting from climate change is a major cause for concern and is expected to exacerbate these risks by increasing the frequency and severity of marine heatwaves and compounding stressors on shellfish. For example, from 2014 to 2016, “the blob” marine heatwave occurred in the northeast Pacific Ocean and caused drastic changes in behavior and populations of forage fish, marine mammals, and sea birds in Washington State (Bond et al., 2015; Cavole et al., 2016; Oliver et al., 2018; Tate et al., 2021). This marine heatwave, along with the already occurring increase in global sea surface temperatures (Cael et al., 2024), has contributed to the poor condition of the animals from plankton biomass to fishes and local birds and mammals (Rogers et al., 2020). At the same time, warming waters favor the proliferation of bacteria and viruses (Estrada et al., 2021; Okon et al., 2023), such as the ostreid herpesvirus (OsHV-1), which can cause devastating losses in densely farmed oyster populations (Renault et al., 2014), prompting efforts to develop disease resistant oyster lines for commercial production (Liu et al., 2023). As selective breeding programs develop oyster lines with disease tolerance, it is unclear whether there are tradeoffs with performance and tolerance to other stressors (Dégremont et al., 2015). Together, these factors highlight the urgent need to understand how oysters cope with multiple, interacting stressors and develop strategies to increase performance under variable environments.

Environmental memory is one mechanism by which organisms may improve stress tolerance within and across generations (Byrne et al., 2020; Kim et al., 2024; Lee et al., 2020; Liu et al., 2023; Ross et al., 2016), demonstrated, for example, in plants (Zhang et al., 2025), corals (Putnam and Gates, 2015), urchins (Chamorro et al., 2023), and copepods (Lee et al., 2022). In oysters, emerging evidence indicates that prior exposure to environmental stress can induce molecular and physiological changes including immune activation, metabolic adjustments, and epigenetic regulation that have the potential to enhance stress tolerance. For example, stress priming has been shown to alter gene expression (Lafont et al., 2020), and upregulation of receptors for hormones and neurotransmitters (Li et al., 2022), and increase resilience (Pereira et al., 2020). Although priming does seem to have physiological effects, it remains uncertain whether priming can meaningfully improve performance in aquaculture environments on time scales that are relevant for commercial production (de Kantzow et al., 2023; Lafont et al., 2017). It also remains unclear whether stress priming can effectively alter offspring performance in cross-generational applications. In one example, (Song et al., 2025) tested for transgenerational immune priming (TGIP) in *M. gigas* offspring, observing bacteria-mediated immunological priming that improved performance in offspring. There is a need for further testing of TGIP effects and evidence is lacking on the effects of stress priming on tolerance to multiple stressors or tolerance to different stressors than the priming experience (i.e., cross-priming). Therefore, there is a need to test the capacity for immune priming in oysters and examine potential implications for broader stress resilience.

Given the multi-stressor environments oysters are exposed to, there is also a need to understand interactions between immune responses and other stress response pathways (i.e., thermal stress response). Oysters possess an innate immune system capable of mounting responses to microbial challenges, despite lacking adaptive immunity (Li et al., 2026). Immune priming in oysters has been induced experimentally using Poly(I:C), a synthetic double-stranded RNA (dsRNA) that mimics viral infection (Lafont et al., 2020). Poly(I:C) activates toll-like receptor 3 and downstream signaling pathways, triggering cytokine production and other immune responses (Doukas et al., 1994). Importantly, Poly(I:C) exposure has been shown to protect oysters against subsequent pathogen challenge (de Lorgeril et al., 2018; Green and Montagnani, 2013; Lafont et al., 2017; Martins, 2020). Therefore, the capacity for synthetic immune exposures to elicit environmental memory within and across generations should be evaluated.

Oysters survive in highly dynamic environments, where their immune responses interact with complex and interacting stressors, raising questions on how immune priming may impact response to other environmental stressors (e.g., thermal stress). Oyster expanded and polymorphic gene families have diverged to create structural diversity, enabling certain receptors to have multiple roles in responding to abiotic stress and pathogens (Guo et al., 2015).

Specifically, both heat and immune stressors create a positive feedback loop of oxidative damage and inflammation in tissues that require similar cellular responses (Rahman et al., 2019; Wang et al., 2012). Under chronic heat stress, for example, persistent low-grade immune activation is maladaptive, reducing overall fitness (Cantet et al., 2021). Short, controlled immune priming using heat stress in aquaculture, where the stress exposure is brief, has improved subsequent pathogen resistance (Cantet et al., 2021).

There is a gap in knowledge on whether immune priming can persist across generations to impact offspring stress tolerance. If immune challenge in broodstock enhances the resilience of their offspring to additional stressors, this could provide a powerful mechanism linking parental experience to offspring survival in aquaculture and natural populations. To address these questions, adult *M. gigas* were exposed to an immune challenge using Poly(I:C) immersion immediately prior to strip spawning and the offspring were reared to the seed stage. Offspring were then subjected to thermal stress experiments to evaluate growth, survival, and metabolic response. We hypothesized that offspring of Poly(I:C)-primed parents would show improved growth, survival, and metabolic performance under thermal stress relative to offspring of unprimed parents. We further tested whether parental Poly(I:C) treatment induced changes in offspring thermal stress response and immunity by characterizing expression of two heat shock proteins, HSP70 and HSP90 (ATP-dependent molecular chaperone proteins), and a gene involved in innate immunity, viperin (Virus Inhibitory Protein, Endoplasmic Reticulum-associated, Interferon-inducible) under acute thermal stress. By integrating immune priming and thermal stress experiments, this study tests whether parental immune challenge influences offspring performance and thermal tolerance as a result of immune priming. Understanding parental effects on oyster stress tolerance is essential for predicting future oyster survival and developing aquaculture strategies that enhance resilience under intensifying thermal and pathogenic stress.

## Methods

### A. Broodstock exposure and fertilization

To test effects of parental immune priming on offspring performance, *Crassostrea/Magallana gigas* broodstock (50-80 mm; 3-4 years old) were exposed to Poly(I:C) immersion prior to strip spawning on October 7, 2024. Broodstock were collected from Sequim Bay, WA and provided by Jamestown S’Klallam Point Whitney Shellfish Hatchery (Brinnon, WA). Broodstock were randomly assigned to either control or Poly(I:C) treatments. Poly(I:C) is a non-infectious elicitor and viral mimic in the form of synthetic double stranded RNA (dsRNA) that has been widely used to induce immune responses in many organisms (Hafner et al. 2013, Lafont et al. 2017). Poly(I:C) exposure has been shown to elicit protective immune responses in oysters in response to disease and infections (de Lorgeril et al., 2018; Green and Montagnani, 2013; Lafont et al., 2017; Martins, 2020). The most common application of Poly(I:C) is direct injection in anesthetized animals, but (Lafont et al., 2017) found that immersion in a 0.38-0.75 µg mL^-1^ Poly(I:C) seawater solution provided equivalent protective effects to injection. The immersion method was applied in this study to conduct Poly(I:C) exposures on broodstock. Oysters (n=30-40 per treatment) were held for 2 h in two separate 100 L bins filled with ambient seawater with constant bubbling (n=1 bin per treatment). The bin with treated oysters included the addition of Poly(I:C) adjusted to a concentration of 0.75 µg mL^-1^ following (Lafont et al., 2017).

Following exposure, oysters were strip spawned following the Jamestown S’Klallam Seafood spawning protocols. Oysters were opened and sexed and selected for high-quality gametes (i.e., large, round eggs and concentrated sperm with low debris). Female gonad tissue was isolated from selected females and added to a plastic bag and gently rubbed to homogenize for each treatment separately. Gonad tissue was poured onto a 20 µm screen and rinsed with 1 µm filtered seawater (FSW) into a 3.7 L bucket held at 25-26°C to hydrate for 20-30 min. Sperm was pooled from all males for each treatment separately and added to a glass dish with 1 µm FSW. Motility was observed using a dissecting microscope. Following egg hydration, approx. 10 mL of sperm was added from the respective treatment to fertilize eggs. Fertilized eggs were then added to a 5,000 L tank with FSW (n=1 tank per treatment) held at 25-26°C and 8.25-8.27 pH (National Bureau of Standards (NBS) scale). In total, the control group included gametes from 12 males and 14 females, and the treated group included gametes from 14 males and 15 females.

### B. Larval rearing

Larvae were reared following the Jamestown S’Klallam Seafood larval rearing protocols conducted by hatchery staff. From 0-3 days post fertilization (dpf), larvae were reared in 5,000 L tanks (n=1 per treatment) at 25-30°C and 8.2-8.4 pH buffered with calcium carbonate with 20 µm banjo filters and 7 L min^-1^ flow. EDTA (1 g per 1,000 L) was added daily to prevent microbial growth. Larvae were fed daily starting at 1 dpf with a mixture of *Pavlova* spp. and *Chaetoceros calcitrans* (fluorescence 5-10; FluoroSense Handheld Fluorometer, Turner Designs, San Jose, CA, USA). At 3 dpf, larval tanks were dropped onto 60 µm mesh sieves. Larval density was estimated as 54 million larvae in the control group and 29 million larvae in the treated group. Larvae were then redistributed into 1,000 L tanks at 18 million per tank for control larvae (n=3 tanks) and 14.5 million per tank for treated larvae (n=2 tanks). 1,000 L tanks were equipped with 60 µm banjo filters with 1.5-2 L min^-1^ flow with daily EDTA addition and feeding through 8 dpf (fluorescence 10-15) at 25-30°C and 8.2-8.4 pH. Environmental conditions are displayed in **Table 1**.

**Table 1.** Environmental conditions (mean ± st. dev.) during larval rearing at the Point Whitney Shellfish Hatchery. Temperature, pH, and salinity were measured using an Orion Star A325 multi-meter. pH was recorded in National Bureau of Standards (NBS) units. Feeding fluorescence during larval rearing was measured using a FluoroSense Fluorometer.

| Time Period | Temperature (°C) | pH (NBS) | Salinity (psu) | Feeding | Tank Volume (L) |
| --- | --- | --- | --- | --- | --- |
| 0-3 dpf | 26.7 ± 2.78 | 8.11±0.23 | 30.9 ± 0.06 | Daily (fluorescence = 5-10) | 5000 L (n=2) |
| 4-8 dpf | 28.5 ± 1.17 | 8.16 ± 0.18 | 31.0 ± 0.04 | Daily (fluorescence = 10-15) | 1000 L (n=5) |
| 9-14 dpf | 28.4 ± 0.35 | 8.15 ± 0.05 | NA | Daily (fluorescence = 20-40) | 1000 L (n=5) |
| Seed grow out at hatchery | 7.61 ± 0.23 | 7.70 ± 0.05 | 23.8 ± 0.46 | Continuous, ambient seawater | 100 L upwellers (n=4) |
| Seed holding at UW | 11.1 ± 0.35 | 7.92 ± 0.10 | 26.9 ± 1.07 | Twice weekly (algal paste) | 115 L (n=2) |

On 8 dpf, larvae were dropped onto a series of sieves (180-30 µm sieves) to remove debris and remove small larvae. Larvae retained on the 105 µm sieves were kept and density was estimated by weighing larvae. There were 50 million larvae in the control group (survival from 3 dpf = approx. 91%) and 27 million in the treated group (survival from 3 dpf = ∼93%). 1,000 L tanks were then cleaned, refilled, and fitted with 105 µm banjo filters. Larvae were added at a density of 13.5 million per tank in the treated group (n=2 tanks) and 16.5 million per tank in the control group (n=3 tanks) with larvae allocated to tanks randomly. Tanks were maintained at 28-30°C and pH 8.1-8.3 with daily EDTA addition and increased feeding (fluorescence 20-40) with flow set at 2-2.4 L min^-1^.

At 14 dpf, larvae were again dropped onto 240 µm sieves. Larvae were well developed with high activity, pigmentation, and feet and eye spots. Larval density was estimated at 25.8 million larvae in the treated group (>95% survival) and 51 million in the control group (∼100% survival). Larvae were wrapped in filter paper and prepared for shipment to the Jamestown Seafood hatchery facility in Kona, Hawaii for setting on 21 October 2024.

### C. Larval setting and transport to UW

Larvae were set at the Jamestown Seafood hatchery facility in Kona, Hawaii following commercial protocols and conducted by hatchery staff. Eyed prediveliger larvae were isolated on screens (236-255 µm) and refrigerated on a damp cloth in a plastic bag for 24 h. Larvae were then stocked into 60 x 90 cm downwelling settlement trays using crushed oyster shell (graded 180-500 µm) as substrate. Stocking density averaged 2 million larvae per tray. Treatment groups were set separately. Seawater was mixed to include surface (warm) and deep (cold) water at 23°C with sand filtration. During setting, larvae were fed continuously with a mixture of *Tetraselmis*, *Tisochrysis*, and *Chaetoceros* spp. Larvae were screened off of the substrate after 8 days in settlement trays. Setting rates were estimated to be high and similar for both treatments (20-25%). Set seed were then shipped back to the Point Whitney hatchery (Brinnon, WA) on 22 November 2024. Seed were then allocated to n=2 indoor upwellers per treatment (0.9 L seed per tank in control groups, 0.75 L seed per tank in treated groups) supplied with ambient seawater (**Table 1**). Hobo TidBit temperature loggers were added to n=2 tanks to monitor temperature every 15 min.

Seed were transported to the University of Washington for experiments on 26 February 2025 and 22 March 2025. Seed were held at the University of Washington in two 115 L tanks held at 14°C and filled with water adjusted with Instant Ocean salts (Instant Ocean, Blacksburg, VA) and equipped with constant bubbling and recirculating pumps. Seed were fed twice weekly (Reed Mariculture Shellfish Diet 1800) and water was changed weekly.

### D. Seed growth

Images were collected throughout seed rearing at the Point Whitney Shellfish Hatchery to monitor seed growth. Images were collected at the following time points: 4 December 2024 (59 dpf), 7 January 2025 (93 dpf), 25 February 2025 (123 dpf), and 30 May 2025 (236 dpf). Shell length (mm) was measured by selecting 100 random oysters from each image for maximum shell length (i.e., hinge to shell edge along the longest axis) measurements using Image J (Rueden et al., 2017).

### E. Survival assays

Survival was assessed by allocating seed to 96-well plates (individual seed in each well) filled with water adjusted to 25 psu and exposed to temperature treatments in benchtop incubators (Vevor, Rancho Cucamonga, CA). Seed were exposed to eight temperature treatments (16°C, 20°C, 21°C, 36°C, 38°C, 40°C, 42°C, and 45°C) for 24 h with seed discarded following each assay. These experiments were conducted over 8 different days spanning two months (February and March 2025) with 16 total plates used (n=96-144 per treatment per temperature). Oysters were 118-216 dpf during resazurin testing with replicates from both treatment groups as well as a control temperature run each day to account for effects of day of measurement.

Mortality was assessed at 5 and 24 h of incubation during survival trials by removing the plates from the incubator and viewing seed under a dissecting microscope and probing with tweezers. If the oyster was observed with an open shell that did not close in response to probing or movement, the oyster was considered dead. Oyster mortality was recorded as a binary response with 0 indicating live and 1 indicating dead oysters.

### F. Resazurin metabolic assays

Resazurin metabolic assays were used to measure metabolic rate in oyster seed across temperatures following the general protocol described in (Huffmyer et al., 2026). Oysters were allocated into 96-well plates (n=1 individual per well) and photographed for shell length measurements (mm). A solution of resazurin (111 µg mL^-1^) was prepared by mixing 0.22% (v/v) resazurin solution (4.76% w/v in deionized water), 0.1% DMSO (v/v), and 1.0% (v/v) antibiotic antimycotic 100x Penn/Strep/Fungizone solution (Cat. SV30079.01, Cytiva, Marlborough, MA) in DI water adjusted to 25 ppt using Instant Ocean salts (Instant Ocean, Blacksburg, VA). Wells with oysters were filled with resazurin solution (180 µL) and read immediately on a plate reader (BioTek FLx800; Agilent, Santa Clara, CA) in fluorescence mode with emission of 530 nm (530/20 nm filter) and excitation of 590 nm (590/20 nm filter) with associated software (BioTek Gen5). Seed were then exposed to control (18-20°C) or high temperature conditions (36°C, 38°C, 40°C, or 42°C) in a benchtop incubator (25 L, Vevor). Trials were conducted over 4 days with 1 control plate and 1 elevated temperature plate run each day with temperatures run in random order across days. There were n=42 control and n=42 treated seed loaded into each plate. Each plate also included n=12 wells of resazurin solution without oysters as blanks. Fluorescence was read every hour for 4 h and measurements were calculated as described below.

### G. Acute stress testing and qPCR

Oyster seed from Poly(I:C) treated parents and control seed were exposed to either an elevated temperature of 44°C (“high”) or held at 10°C for 2 h (“control”) in 24-well plates (n=1 seed per well, n=20 per condition) in a benchtop incubator (25 L, Vevor). Wells were filled with 2 mL of resazurin solution (prepared as described above). Due to sample size limitations, resazurin metabolic responses were not measured in this study. After 2 h, all seed were frozen at −80 °C for subsequent RNA extraction.

Tissues were homogenized in 1.5 mL SafeLock (Eppendorf) microcentrifuge tubes containing 300 uL of DNA/RNA Shield (ZymoResearch), ∼50 uL of 0.5 mm zirconium oxide beads (NextAdvance) and ∼50 uL of 0.15 mm zirconium oxide beads (NextAdvance), using a Bullet Blender 5E Gold + (NextAdvance) with 1.5 mL tube adapters for 3 min at speed 12. The Bullet Blender was pre-cooled, and cooled, using dry ice. After homogenization, tubes were briefly centrifuged at 16,000 g and the supernatants were transferred to new 1.7 mL microfuge tubes. RNA was isolated using the Quick-DNA/RNA MiniPrep Plus Kit (ZymoResearch) according to the manufacturer’s protocol, including the DNase treatment step. RNA was eluted with 50 uL of water. The RNA was quantified with a Qubit 3.0 (Invitrogen) using the Qubit Broad Range RNA Assay (Invitrogen) and stored at −80C.

Reverse transcription was performed using the GoScript Reverse Transcription System (Promega) according to the manufacturer’s protocol. The reaction size was doubled, from 20 uL to 40 uL, to allow for sufficient cDNA for all qPCRs. Each reaction contained 200 ng of total RNA. The optional RNasin was also used. Reactions were set up in a 96-well, standard qPCR plate and sealed with strip caps. The reverse transcriptase components were prepared as a master mix and then distributed to RNA/oligo dT mixtures. All incubations were conducted in a 96-well PTC-200 (MJ Research) thermal cycler using a heated lid. The cDNA was stored at −20C. Quantitative polymerase chain reactions (qPCRs) were run in 20 uL reactions using SsoAdvanced Universal SYBR Green Supermix (BioRad), in triplicate, in low-profile, white 96-well plates (USA Scientific) sealed with TemPlate TR Select Optical Film (USA Scientific) in a CFX Connect (Bio-Rad) or CFX96 (Bio-Rad) real-time thermal cycler. Reactions were prepared as master mixes and distributed to wells, followed by addition of cDNA or water (no template control [NTCs]). A single reaction contained the following: 1 uL of cDNA (or water for NTCs), 0.25 uM each of forward and reverse primers, 10 uL of SsoAdvanced Universal SYBR Green Supermix, and water to a final reaction volume of 20 uL. Only one primer set was run per plate. Thermal cycling conditions were as follows: 1 cycle of 95°C for 30 s; 40 cycles of 95°C for 3 s, 60°C for 5 s; dissociation analysis from 65°C to 95°C in 0.5°C increments, each held for 5 s. Baselines and thresholds were autocalculated by the CFX Maestro (Bio-Rad) software. The dissociation curves for all samples and NTCs were evaluated on all plates to ensure there was no contamination and/or spurious amplification.

### H. Data analysis

All data were analyzed using R Statistical Programming (v4.4.2; (R Core Team, 2022)). Growth was analyzed using a linear model with length (mm; log10 transformed) as the response variable and parental treatment, time (dpf), and their interaction as main effects. Significance of main effects was assessed with a Type II ANOVA test with Satterthwaite’s approximation in the car package (Fox and Weisberg, 2018). Post hoc tests were conducted using estimated marginal means in the *emmeans* package (Lenth, 2018). Model assumptions were evaluated by examining diagnostic plots of residuals versus fitted values and scale-location plots to assess homogeneity of variance, and normality of residuals was assessed using quantile–quantile plots.

Survival under thermal stress (24 h) was analyzed using a binomial logistic regression with mortality (0=live, 1=dead) as the response variable and temperature, parental treatment, and their interaction as main effects. Tank and batch were included as random intercepts.

Significance was assessed using Type II Wald Chi Square ANOVA tests in the *car* package (Fox and Weisberg, 2018). Post hoc tests were also performed using estimated marginal means in the *emmeans* package (Lenth, 2018). Model predictions of survival were plotted using the *ggeffects* package (Lüdecke, 2018). Model assumptions were evaluated using simulation-based residual diagnostics implemented in the DHARMa package (Hartig, 2024), including tests for overdispersion and zero inflation.

Metabolic rates were calculated from resazurin fluorescence as described in (Huffmyer et al., 2026). Briefly, fluorescence measurements were first normalized to the initial fluorescence value to generate fold change in fluorescence and then corrected by subtracting mean normalized fluorescence values of blanks. Corrected fluorescence values were then normalized to size by dividing by shell length, producing metabolic rates expressed as fold change in fluorescence mm^-1^. Total metabolic activity was calculated for each oyster as Area Under the Curve (AUC) using trapezoidal integration in the *pracma* package (Borchers, 2023). Total metabolic activity (AUC) was then analyzed using a linear mixed effect model with log-transformed AUC as the response and temperature, parental treatment, and their interaction as main effects. Tank and plate nested within date was included as a random intercept. Significance was assessed using Type II ANOVA tests with Satterthwaite’s approximation with post hoc tests performed using estimated marginal means in the *emmeans* package (Lenth, 2018). Model predictions of metabolic activity were plotted using the *ggeffects* package (Lüdecke, 2018). Model assumptions were evaluated by examining diagnostic plots of residuals versus fitted values and scale-location plots to assess homogeneity of variance, and normality of residuals was assessed using quantile–quantile plots.

We selected two heat shock proteins, HSP70 and HSP90 (ATP-dependent molecular chaperone proteins), and a gene involved in innate immunity, viperin (Virus Inhibitory Protein, Endoplasmic Reticulum-associated, Interferon-inducible), for quantitative polymerase chain reaction (qPCR) analysis to test whether Poly(I:C) parental treatment induced changes in organism thermal stress response and immunity. We selected GAPDH (Glyceraldehyde 3-phosphate dehydrogenase) as a normalizing gene due to its stable Cq values across all samples assayed. Technical replicates with Cq standard deviations > 0.5 were excluded from analysis. The ΔCq method was used for normalization, using the mean Cq of technical replicates for each target and GAPDH. (ΔCq = mean Cq, target - Cq, GAPDH).

ΔCq values for HSP70, HSP90, and viperin were analyzed using two-way ANOVA tests with acute stress treatment (i.e., control or high temperature) as main effects. Assumptions of residual normality and homogeneity of variance were verified using quantile-quantile plots and scale-location plots, respectively. Post hoc tests were performed using estimated marginal means in the *emmeans* package (Lenth, 2018). Due to variation in sample size between groups, we performed a downsampling analysis to determine the robustness of significance results to unequal sample size. For each gene, minimum sample size was identified with the lowest sample size occurring in the Poly(I:C) treated - high temperature group (HSP70 n=18, HSP90 and viperin n=8). Data were then randomly subsampled without replacement to the minimum n from treatment group and the ANOVA model was re-run for 1,000 iterations. For each ANOVA model term, we report the proportion of iterations in which the term was significant at P<0.05. Final results are presented from the full ANOVA model with all available observations. Log2 fold change (log2 FC) values were calculated from ΔΔCq as log2 FC = −ΔΔCq, where ΔΔCq = ΔCq_sample − mean(ΔCq_reference), with the control Poly(I:C) - control temperature group used as the reference. Individual sample log2 FC values were plotted with the mean log2 FC of the reference group equal to 0. Cohen’s D effect sizes were calculated for the effect of Poly(I:C) treatment on ΔCq values of each gene.

## Results

### A. Growth rates were modulated by parental treatment

Shell length in seed increased across development and was modulated by parental treatment (P _Treatment x Time_ <0.001; **Table 2**). At 59-93 dpf, seed from control parents (“control”) were larger (control 4.43 ± 0.06 mm; treated 3.88 ± 0.04 mm; post hoc P < 0.001), but by 236 dpf, seed from treated parents (“treated”) were larger (control 9.14 ± 0.14 mm; treated 9.74 ± 0.16 mm; post hoc P = 0.013; **Fig 1**).

**Fig 1.**
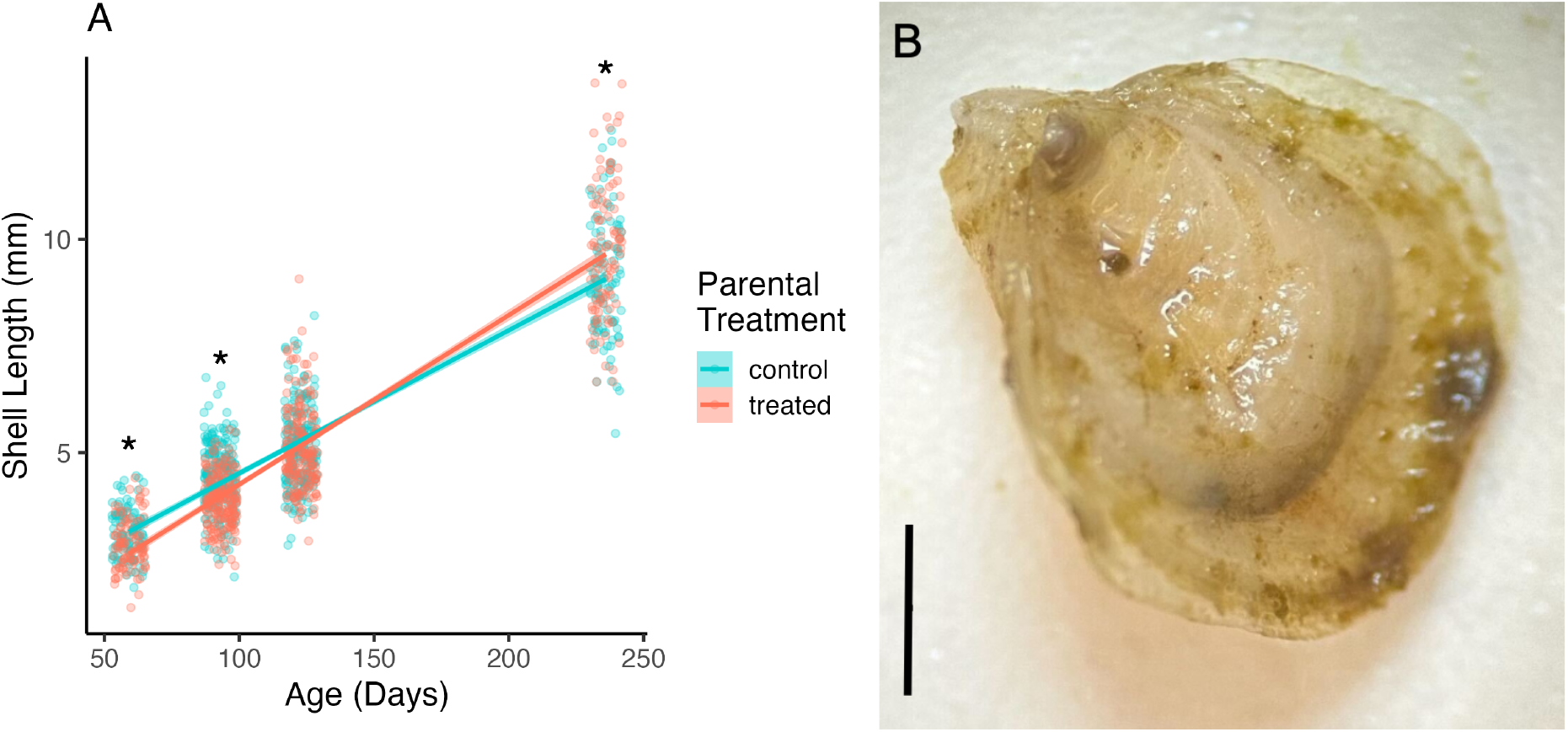
(A) Growth of seed from control and treated parental groups (blue and red, respectively) from December 2024 to June 2025 (59-236 dpf). Line indicates linear model of shell length over time with individual data points overlaid. Asterisks indicate significant differences (P < 0.05) between control and treated seed. (B) Image of an example specimen of a Pacific Oyster seed in this study with a scale bar representing 1 mm.

**Table 2.**
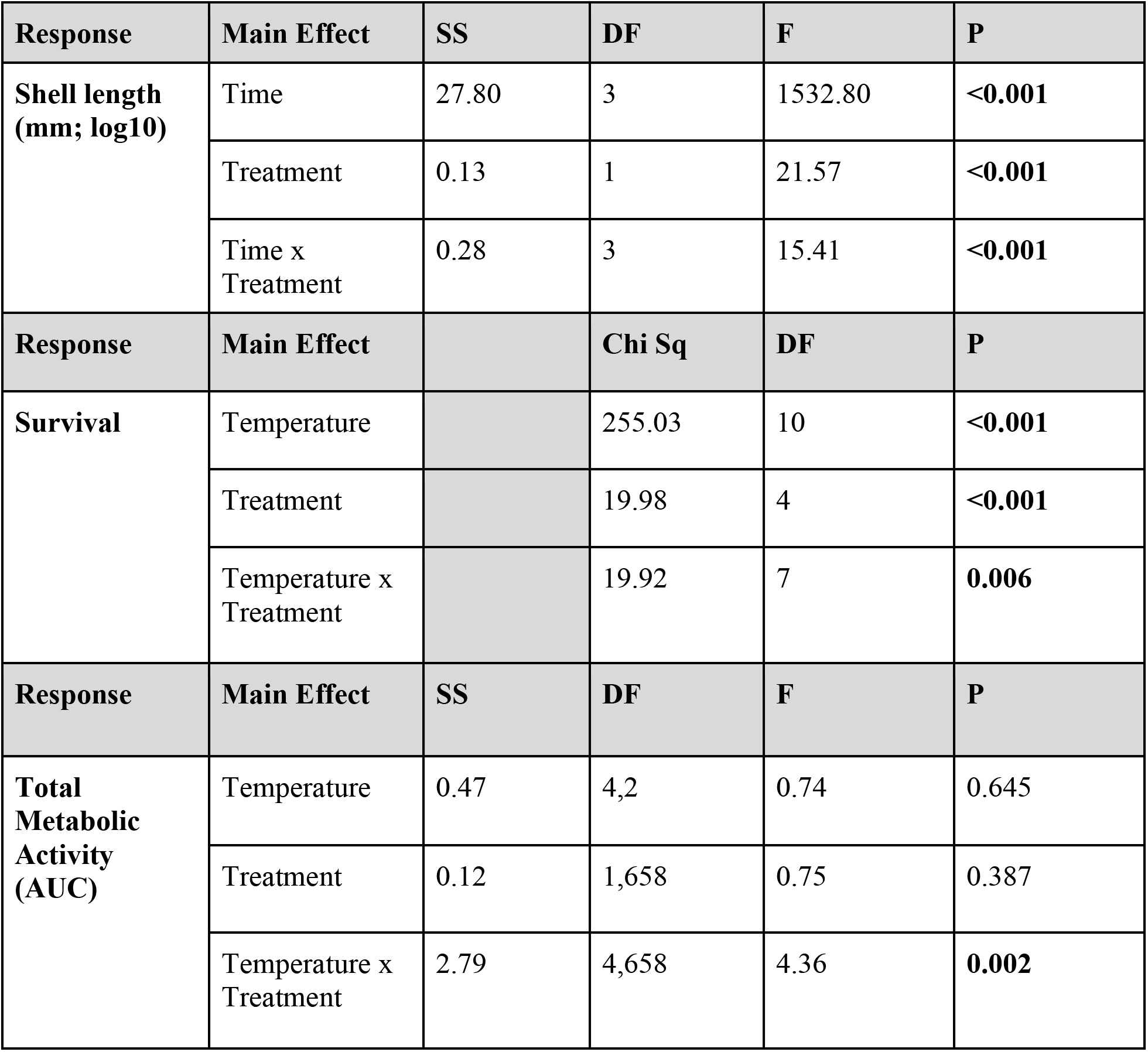
Statistical analysis of responses of oyster seed. Growth was analyzed using a linear model with significance determined using Type II ANOVA. Total metabolic activity (area under the curve; AUC) was analyzed using a linear mixed effect model with significance determined by Type II ANOVA with Satterthwaite’s approximation. Survival was analyzed using a generalized mixed model (family = binomial, link = logit) with significance determined by Type II Wald Chi Square ANOVA. SS = sum of squares; DF = degrees of freedom.

### B. Differential survival under thermal stress

Thermal tolerance was measured as survival during temperature exposure (16-45°C) in the lab with survival observed at 5 and 24 h of exposure. Mortality increased with temperature with significant effects of parental treatment on survival at elevated temperatures (P _Temperature x Treatment_ = 0.006; **Table 2**; **Fig 2**). Specifically, thermal tolerance (i.e., lower mortality) was higher for treated seed at 40°C (post hoc P = 0.007), but tolerance was lower (i.e., higher mortality) in treated seed at 42°C compared to control (post hoc P = 0.018) with no significant differences at other temperatures (post hoc P > 0.05; **Fig 2**). The greatest difference in thermal tolerance was observed at 40°C with probability of mortality 35% lower in treated seed. At 42°C, however, mortality was 17% higher in treated seed compared to control (**Fig 2**).

**Fig 2.**
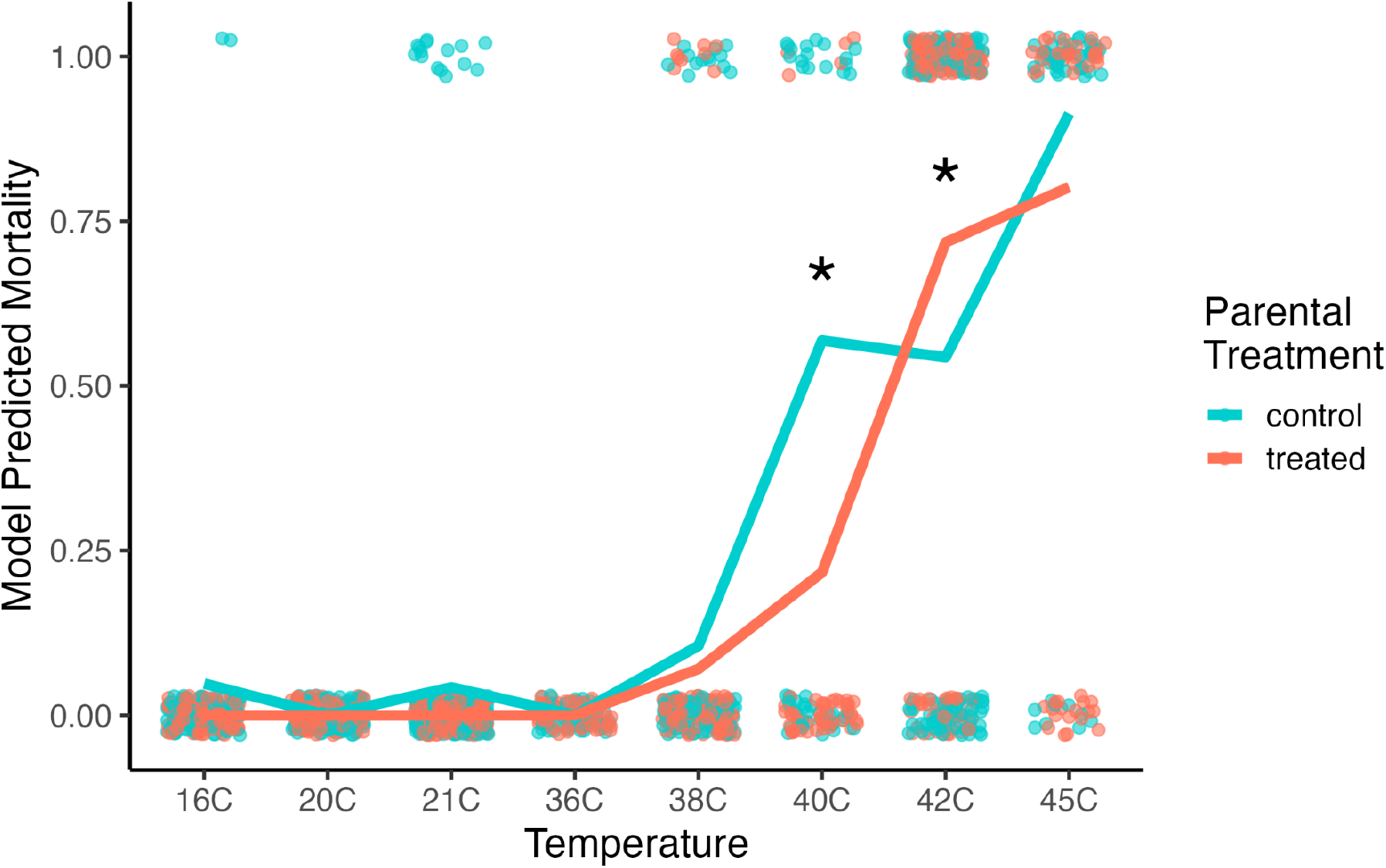
Total mortality in control (blue) and treated (red) seed after 24 h of exposure to temperature stress tests ranging from 16-45°C. Lines show logistic regression model predicted mortality with individual points showing oysters observed as dead (1) and alive (0). Asterisks indicate significant (P < 0.05) post hoc comparisons between treatments at each temperature.

### C. Metabolic differences in temperature response between treatments

Resazurin metabolic assays (**Fig 3**) were conducted to examine differences in metabolic response to temperature treatments ranging from 20-42°C in the lab. Total metabolic activity is represented as area under the curve (AUC). Metabolic activity of seed across temperature was modulated by parental treatment (P _Temperature x Treatment_ = 0.002). Specifically, metabolic activity was significantly higher (43%) in treated seed compared to control seed at 36°C (post hoc P = 0.003). At 40°C, however, metabolic activity was (30%) higher in control seed (post hoc P = 0.019; **Fig 3**). There was no difference in metabolic activity at any other temperature (post hoc P > 0.05 for all comparisons).

**Fig 3.**
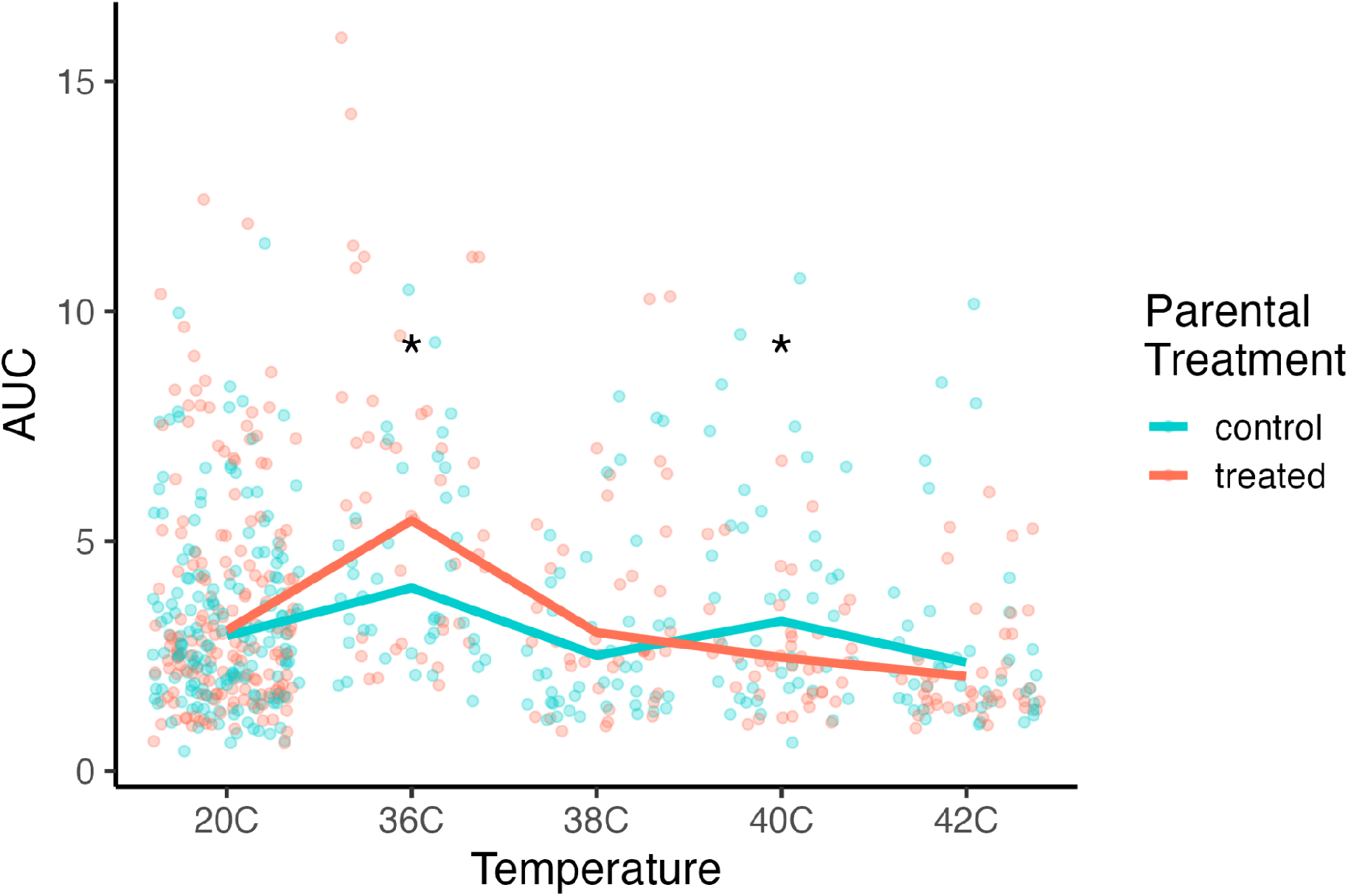
Total metabolic activity (area under the curve; AUC) measured using the resazurin assay. Lines show linear regression model predicted AUC with individual points for control (blue) and treated (red) seed (n = 42 control and n = 42 treated seed per plate). Asterisks indicate post hoc comparison P < 0.05.

### D. Variation in gene expression under acute stress exposure

Expression of HSP70, a heat shock protein, was significantly higher under acute 44°C temperature conditions (P < 0.001; **Fig 4A**; **Table 3**). Poly(I:C) parental treatment had a large and significant effect on expression (P < 0.001) with elevated expression in treated seed at both 10°C (post hoc P < 0.001; *d* = −1.28; log2 FC = 2.22) and 44°C temperature (post hoc P = 0.021; *d* = −0.82; log2 FC = 6.20; **Fig 4A**; **Table 3**). Expression fold change values and group sample sizes are presented in **Table 4**. Note that negative Cohen’s D effect size values (*d*) indicate a lower ΔCq and therefore reflect the observed higher gene expression in the Poly(I:C) treated group.

**Fig 4.**
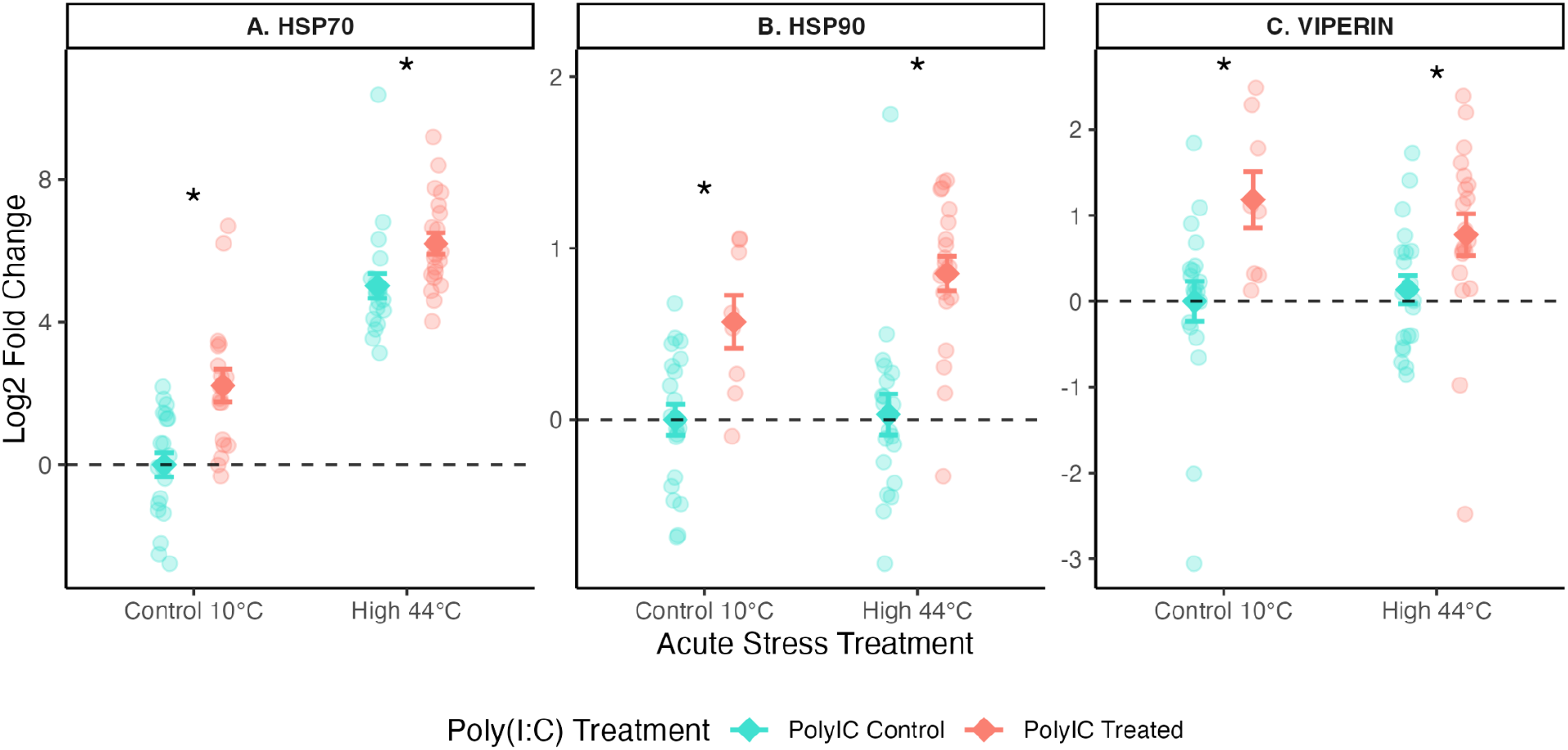
Log2 fold change (FC) in gene expression relative to the control Poly(I:C) - control temperature group for HSP70, HSP90, and viperin genes at control (10°C) and high temperature (44°C) acute stress treatments. Individual data points represent individual sample log2 FC values; diamonds indicate group means ± standard error of mean. Cyan = Poly(I:C)-control; salmon = Poly(I:C)-treated. Dashed line indicates no change in expression (log2 FC = 0; equivalent to 1-fold change). Asterisks indicate significant (P < 0.05) post hoc comparison between Poly(I:C) treatments at the respective temperature.

**Table 3.** Statistical analysis of ΔCq values of heat shock proteins (HSP70, HSP90) and an immune gene (viperin) analyzed by qPCR in oyster seed. ΔCq values were analyzed using Type II ANOVA with acute stress treatment and PolyIC treatment as main effects. SS = sum of squares; DF = degrees of freedom. Proportion P < 0.05 in downsampling indicates the proportion of tests out of 1,000 iterations in which the model term was significant at P < 0.05 when downsampling to minimum group sample size without replacement. Bold indicates P < 0.05 with robust downsampling results.

| Gene | Main Effect | SS | DF | F | P | Proportion $P < 0.05$ down-sampling |
| --- | --- | --- | --- | --- | --- | --- |
| **HSP70** | Acute stress treatment | 396.4 | 1 | 156.71 | **<0.001** | **100%** |
|  | PolyIC treatment | 63.6 | 1 | 25.13 | **<0.001** | **100%** |
|  | Acute stress x PolyIC | 5.2 | 1 | 2.07 | 0.155 | 0.4% |
| **HSP90** | Acute stress treatment | 0.23 | 1 | 1.09 | 0.300 | 6.3% |
|  | PolyIC treatment | 9.43 | 1 | 44.01 | **<0.001** | **99.8%** |
|  | Acute stress x PolyIC | 0.23 | 1 | 1.07 | 0.304 | 8.2% |
| **viperin** | Acute stress treatment | 0.06 | 1 | 0.06 | 0.801 | 1.2% |
|  | PolyIC treatment | 11.16 | 1 | 11.97 | **0.001** | **80%** |
|  | Acute stress x PolyIC | 1.08 | 1 | 1.16 | 0.286 | 1.9% |

**Table 4.** Fold change (FC) and log2 FC in gene expression relative to the control Poly(I:C) - control temperature group for HSP70, HSP90, and viperin genes at control (10°C) and high temperature (44°C) acute stress treatments. SE indicates standard error of mean.

| Gene | Poly(I:C) Treatment | Stress Treatment | n | Mean log2 FC | SE log2 FC | Mean FC | SE FC |
| --- | --- | --- | --- | --- | --- | --- | --- |
| **HSP70** | Control | 10°C Control | 20 | 0.00 | 0.34 | 1.00 | 0.24 |
|  | Treated | 10°C Control | 18 | 2.22 | 0.46 | 4.66 | 1.48 |
|  | Control | 44°C High | 20 | 5.02 | 0.34 | 32.36 | 7.70 |
|  | Treated | 44°C High | 20 | 6.20 | 0.30 | 73.52 | 15.29 |
| **HSP90** | Control | 10°C Control | 20 | 0.00 | 0.09 | 1.00 | 0.06 |
|  | Treated | 10°C Control | 8 | 0.57 | 1.16 | 1.49 | 0.16 |
|  | Control | 44°C High | 20 | 0.03 | 0.12 | 1.02 | 0.09 |
|  | Treated | 44°C High | 20 | 0.85 | 0.10 | 1.81 | 0.13 |
| **viperin** | Control | 10°C Control | 20 | 0.00 | 0.23 | 1.00 | 0.16 |
|  | Treated | 10°C Control | 8 | 1.18 | 0.33 | 2.27 | 0.51 |
|  | Control | 44°C High | 20 | 0.14 | 0.17 | 1.10 | 0.13 |
|  | Treated | 44°C High | 20 | 0.78 | 0.24 | 1.71 | 0.29 |

Similarly, there was a large and significant effect of Poly(I:C) parental treatment on expression of another heat shock protein, HSP90 (P < 0.001) but there was no change in expression due to acute stress temperature alone (P = 0.300; **Fig 4B**; **Table 3**). HSP90 expression was higher in Poly(I:C) treated seed at both 10°C (post hoc P = 0.005; *d* = −1.38) and 44°C conditions (post hoc P < 0.001; *d* = −1.67; **Fig 4B**).

Poly(I:C) treatment significantly influenced expression of viperin (P = 0.001; **Table 3**) with a large effect at 10°C (*d* = −1.17; log2 FC = 1.18) and a medium effect at 44°C (*d* = −0.68; log2 FC = 0.78; **Fig 4C**; **Table 4**). Acute stress temperature did not affect viperin expression (P = 0.801). Seed from Poly(I:C) treated parents exhibited higher expression of viperin at 10°C (post hoc P=0.005) and 44°C conditions (post hoc P = 0.041; **Fig 4C**).

For all genes, results were robust to unequal sample sizes (**Table 4**), with the effect of Poly(I:C) treatment remaining significant in 100%, 99.8%, and 80% of downsampling iterations for HSP70, HSP90, and viperin, respectively (**Table 3**). The lower iteration significance rate for viperin is consistent with the observed moderate effect size (*d* = −0.684). The Poly(I:C) effect for viperin was significant in the full dataset (P = 0.001; **Table 3**) but showed greater sensitivity to sample size variation in downsampling (80% of iterations significant). This suggests that while the effect is robust, unequal sample size across treatment groups may contribute to the strength of the observed Poly(I:C) effect on viperin expression.

## Discussion

This study demonstrates that immune priming in *Crassostrea/Magallana gigas* (Pacific oyster) broodstock with a viral mimic, Poly(I:C), reprogrammed offspring physiology and altered cross-generational thermal tolerance. Offspring from immune-challenged parents exhibited coordinated shifts in molecular, metabolic, and organismal responses. Specifically, parental immune priming resulted in frontloaded expression of stress and immune response genes, altered metabolic response to heat stress, modified growth trajectories, and changes in thermal tolerance in offspring. These effects were strongly dependent on temperature, suggesting that cross-generational priming effects are highly context dependent. These results provide evidence that parental immune priming can reshape offspring physiology in Pacific oysters.

While there is potential for priming protocols to positively impact aquaculture resilience, additional research is required to characterize the mechanisms of these effects, potential physiological trade-offs, and environmental limits prior to implementation in shellfish breeding and hatchery operations.

### A. Parental immune priming reprogrammed offspring physiology and altered stress phenotypes

Immune priming in *Crassostrea/Magallana gigas* broodstock with a viral mimic, Poly(I:C), reprogrammed offspring physiology through constitutive upregulation of stress response genes and increased metabolic flexibility. Elevated expression of heat shock proteins (HSP70, HSP90) and an immune gene, viperin, in this study suggest that parental immune priming exposure may result in constitutive upregulation of stress-response pathways and innate immunity in offspring to manage oxidative and inflammatory stress. In oysters, which rely solely on innate immunity, parental effects and non-genetic inheritance may provide the mechanisms for cross-generational responses to stressors experienced by the parents (Gavery and Roberts, 2010; Li et al., 2017). Previous work has also demonstrated that viperin expression is upregulated following Poly(I:C) exposure (Delisle et al., 2023; Green et al., 2015, 2014) and can persist over longer time periods (Lafont et al., 2017), validating that parental treatment elicited immune responses and now demonstrate a cross-generational response that was persistent >7 months after parental exposure.

Rather than exclusively enhancing antiviral defenses, parental immune priming appeared to alter the baseline stress-response state of offspring, activating pathways that may contribute to differences in metabolic regulation during thermal stress. Upregulation of heat shock genes (HSP70, HSP90) suggests that immune priming may have broadly enhanced cellular stress-response pathways that overlap with heat stress responses including reactive oxygen species (ROS) management, stress-response signaling, and immune defenses (Cantet et al., 2021; Lang et al., 2009). These changes in stress-response pathways were accompanied by altered metabolic responses during thermal challenge, suggesting that parental immune exposure influenced how offspring allocate energy under stress.

Offspring from immune-treated parents displayed elevated metabolic rates at moderate levels of thermal stress (36°C) compared to controls, but this effect reversed at elevated temperatures with lower metabolism at higher temperatures (40°C). This pattern suggests enhanced metabolic flexibility, as offspring from immune-primed parents exhibited a greater capacity to modulate metabolic rate across temperatures (Goodpaster and Sparks, 2017).

Increased metabolic rates under moderate stress may support maintenance of cellular stress responses, while reductions in metabolic rate at extreme temperatures may represent an advantageous metabolic depression response to conserve energy (Pörtner, 2008; Sokolova, 2021), which has been associated with higher survival in oysters (Huffmyer et al., 2026).

The relationship between metabolic rate and survival was context-dependent in this study. Immune-primed offspring exhibited reduced metabolic rates and survival at 40°C, but higher survival rates at an even higher temperature of 42°C with no change in metabolic rate. We hypothesize that primed offspring may begin to downregulate metabolism at lower temperatures than non-primed offspring, which sacrifices performance at 40°C while preserving cellular integrity during more extreme conditions, leading to higher survival at 42°C. This nuanced variation in thermal responses suggests that parental immune priming may shift offspring thermal performance curves rather than enhancing thermal tolerance across a broad thermal range. Shifts in offspring performance curves resulting from parental thermal stress exposure have also been demonstrated in the purple sea urchin (*Strongylocentrotus purpuratus*) (Chamorro et al., 2023) and Sydney rock oysters (*Saccostrea glomerata*) (Filippini et al., 2026), with a study in pipefish demonstrating that the effects of parental immune challenges on offspring performance was regulated by thermal environment (Roth and Landis, 2017). Our results further demonstrate that cross-generational impacts of immune priming are strongly driven by thermal environments and additional work should determine environmental limits of priming under marine heat wave scenarios. In organisms that inhabit dynamic environments, parental effects are highly context dependent and additional work is required to determine environmental limits resulting from parental priming exposures.

Physiological trade-offs of parental immune priming were apparent in growth trajectories in which offspring from treated parents grew slower than controls at first but ended up larger by the end of the study. Given that the expression of heat shock and immune genes was upregulated in treated offspring, slower initial growth may represent a physiological cost of frontloading stress response genes (Hamdoun et al., 2003; Sokolova, 2013). This delayed but accelerated growth trajectory suggests a reallocation of energetic investment in early life stages, possibly reflecting tradeoffs between immune preparedness and somatic growth. Previous work in oysters has shown tradeoffs between growth and thermal tolerance (McAfee et al., 2017), though findings in the present study indicate that immune priming in oysters resulted in more complex metabolic tradeoffs. Oyster growth has influence over oyster-virus interactions, as well as other factors including food availability and temperature (Petton et al., 2021). There has been an observed tradeoff between immune response and growth in oysters infected with OsHV-1 where oysters that are exposed to high amounts of food or have a faster growth rate are more susceptible to OsHV-1 than oysters that grow slower (Pernet et al., 2019; Petton et al., 2022).

The mechanisms underlying the cross-generational priming effects observed in this study may involve multiple pathways. We hypothesize that parental immune priming may alter maternal provisioning of the gametes shortly before spawning, which can affect transfer of energy reserves such as lipids (Gibbs et al., 2021; Timmins-Schiffman et al., 2026; Xu et al., 2024), immune- or stress-related transcripts (Green et al., 2016; Lafont et al., 2017), antimicrobial peptides (Sun et al., 2021), and microbial communities (Scanes et al., 2023; Unzueta-Martínez et al., 2022) that may influence offspring physiology. Given that these effects persisted for >7 months after spawning, there may be additional underlying molecular mechanisms such as epigenetic regulation that contribute to upregulation of stress response genes observed in this study (Gawra et al., 2023; Lafont et al., 2020). Because Poly(I:C) exposure occurred within a few hours of spawning in our study, it is also possible that gametes were directly exposed to the viral mimic prior to fertilization. While our experimental design cannot distinguish between gamete effects within the parent prior to spawning and parental effects, the persistence of offspring phenotypes throughout development suggests that parental immune reprogramming was transmitted to offspring through one or more cross-generational mechanisms.

An important limitation of this study is that offspring were initially maintained in one replicate tank per treatment from 0-3 dpf during the earliest stages of development, such that treatment and initial rearing tank effects cannot be statistically distinguished. Several lines of evidence, however, support that Poly(I:C) treatment led to downstream physiological effects on offspring, rather than initial tank conditions. First, offspring exhibited altered expression of viperin, and immune-response gene that has been previously shown to increase following Poly(I:C) exposure in oysters (Delisle et al., 2023; Green et al., 2015, 2014; Lafont et al., 2017), supporting our conclusion that parental treatment elicited expected immune responses. Second, offspring heat shock protein expression varied based on parental treatment, despite similar temperatures in rearing tanks with no exposure to heat stress during early developmental periods. Finally, the effects of parental immune treatment persisted for seven months following parental exposure and were observed across biological levels from gene expression to metabolism and survival, making a transient tank-specific effect less likely. We recommend future experiments use higher tank replication in early life stages to fully separate parental treatment effects from early developmental environment effects.

### B. Implications for oyster resilience and hatchery applications

Any intervention requires extensive additional testing to determine trade-offs, potential impacts, and consistency in direction and magnitude of impacts. The present study demonstrates benefits of parental immune priming at moderate heat stress, but effects are complex and highly temperature dependent, demonstrating that quantifying realized benefits of priming in real-world applications requires further laboratory and field testing. Further, potential tradeoffs between priming and other biological functions such as growth, reproduction, disease susceptibility, and long-term fitness in the natural environment remain untested. The persistence of the documented priming effects beyond early seed stages and their influence on performance are critical next steps for aquaculture applications.

Given the effects of parental immune priming on offspring performance and underlying molecular stress responses, immune priming may have applications to enhance oyster resilience in aquaculture settings as mass mortality events become increasingly common from a combination of environmental factors including heat waves, pathogens, hypoxia, and other stressors (Destoumieux-Garzón et al., 2024). Hatchery-based stress conditioning of broodstock may offer a high-throughput method to improve offspring performance during early life stages and grow out. For example, Poly(I:C) immersion before spawning represents a non-invasive, scalable approach that could be integrated into existing workflows. Integrating priming into hatchery operations and selective breeding programs may offer synergistic benefits of maximizing stress tolerance through both genetic selection and physiological adjustments. These efforts will require unique tailoring to the specific environmental challenges faced by region, farm, and microenvironment. By better understanding and further integrating stress priming into shellfish production workflows, the aquaculture industry may gain additional new strategies for improving oyster stress tolerance and enhancing seafood security.

## Data Availability

All data is openly available on GitHub (https://github.com/RobertsLab/polyIC-larvae) and as a static release (https://github.com/RobertsLab/polyIC-larvae/releases/tag/v2.0).

## Acknowledgements

We gratefully acknowledge the experimental support provided by Matt Henderson and Carrie Speer and the staff at the Jamestown S’Klallam Seafood at the Point Whitney Shellfish Hatchery. We thank Aakriti Vijay at the University of Washington for experimental support and the Roberts Lab at the University of Washington for feedback on data analysis and the manuscript.

## Funding

This work is supported in part by the U.S. Department of Agriculture’s National Institute of Food and Agriculture, project award numbers 2022-70007-38284 and a grant from Washington Sea Grant, University of Washington, pursuant to National Oceanic and Atmospheric Administration Award No. NA14OAR4170078. The views expressed herein are those of the authors and not necessarily reflect the views of funding agencies.

## Notes

https://github.com/RobertsLab/polyIC-larvae/releases/tag/v2.0

### Competing Interest Statement

The authors have declared no competing interest.

### Summary of Updates

This version of the manuscript has been revised to include gene expression data acquired through qPCR. The introduction, methods, results, and discussion have been revised to reflect the new results. Figures and tables were added for gene expression data.

